# The contribution of movement to social network structure and spreading dynamics under simple and complex transmission

**DOI:** 10.1101/2024.02.09.579705

**Authors:** Michael Chimento, Damien R. Farine

**Author notes:** **Materials & Correspondence** Correspondence should be addressed to Michael Chimento.

## Abstract

The structure of social networks fundamentally influences spreading dynamics. In general, the more contact between individuals, the more opportunity there is for the transmission of information or disease to take place. Yet, contact between individuals, and any resulting transmission events, are determined by a combination of spatial (where individuals choose to move) and social rules (who they choose to interact with or learn from). Here we examine the effect of the social-spatial interface on spreading dynamics using a simulation model. We quantify the relative effects of different movement rules (localized, semi-localized, nomadic, and resource-based movement) and social transmission rules (simple transmission, anti-conformity, proportional, conformity, and threshold rules) to both the structure of social networks and spread of a novel behaviour. Localized movement created weakly connected sparse networks, nomadic movement created weakly connected dense networks, and resource-based movement generated strongly connected modular networks. The resulting rate of spreading varied with different combinations of movement and transmission rules, but— importantly—the relative rankings of transmission rules changed when running simulations on static versus dynamic representations of networks. Our results emphasize that individual-level social and spatial behaviours influence emergent network structure, and are of particular consequence for the spread of information under complex transmission rules.

## 1 Introduction

Spreading dynamics, defined as how information, behaviours, or pathogens spread through a population, are fundamental to social evolution. Social behaviours provide the substrate through which information or pathogens can spread, and there is a broad agreement that social structure and interactions play a significant role in population-level consequences (1; 2; 3). The spread of information or pathogens may then affect the balance of costs and benefits of being social (3; 4). This re-balancing of costs and benefits may change social network structure, and thus change future spreading dynamics in a feedback loop. For example, people avoided large social gatherings immediately following an outbreak of Swine flu in Mexico (5), and further modelling found that such rapid re-wiring of social networks led to lower epidemic sizes compared to static networks (6). Similarly, in non-human animals, the social network structure of guppies (*Poecilia reticulata*) was found to change in response to the spread of disease due to active avoidance of infected fish (7). The importance of this feedback means that the impact of social network structure on spread has been well studied, both in terms of the initial spread of disease (8; 9) or information (10; 11) and in terms of how the resulting spreading dynamics can shape social (12; 13; 14) and cultural (15; 16) evolution.

While the dynamics and consequences of spread are becoming well-established, less is known about the factors shaping the structure of animal social networks and how these affect the resulting transmission networks. Networks can be determined by a range of individual and social behaviours. One notable behaviour is movement, which is the most fundamental driver of new contacts. Movement allows organisms to search for resources and avoid threats and has long been recognized as a key link between individuals and higher-level ecological processes (17), such as dispersal (18) and metapopulation connectivity (19). Individuals can move through physical (20; 21) and social (22; 23) space, and the opportunity for pathogens or information to spread is primarily governed by movement decisions of animals (9).

Importantly, movement patterns differ between species, contexts, or life stages, resulting in vastly different social networks. Indeed, spatial movement often accounts for well over half of the structure of social networks (24). Territorial species may exhibit highly localized movement and rarely interact with neighbours, resulting in a lattice-like network (25; 15). Solitary species may move relatively independently resulting in them forming a large number of weak ties (9). In contrast, gregarious species move in synchronous groups resulting in more fragmented, but more cohesive networks (9). For example, a dynamic social network analysis of more sociable Grevy’s zebra (*Equus grevyi*) found modular networks compared with more solitary Asiatic wild ass (*Equus hemionus*) due to differing degrees of synchronous movement (26).

Movement decisions can vary dramatically in both the extent (e.g. migratory vs. non-migratory) and style (e.g. fixed-range vs. open-range), and the resulting effects of temporal changes to networks on spreading have been explored theoretically (27; 28; 29). For example, changing connectivity of social networks from resource-based movement can affect the spreading dynamics of disease (29). However, movement is rarely directly modelled and is instead implicitly encoded into the temporal changes to the network. This has two consequences. First, the contribution of different movement rules on the emergent patterns of social contacts remains opaque. Second, it remains unclear whether networks that summarise observations over time accurately capture the properties of contacts as individuals move in time and over space. These two issues are particularly important when considering the contribution of network structure on transmission dynamics.

It is common when studying networks to condense the temporal dynamics into an averaged static representation of the social contacts. However, the extent to which such representations are suitable for studying information spread is still an open question (30). Disease transmission simulations using dynamic and static representations of the same network data gathered from Verreaux’s sifakas (*Propithecus verreauxi*) found that dynamic simulations could obtain larger estimates of disease outbreak size, but this depended on transmission and recovery parameters (31). Another simulation study found that the predicted dynamics of static networks are likely to match those of dynamic networks if the transmission probability is constant across contacts. By contrast, if the transmission probability varies according to, for example, the type of contact, then the two representations will generate different predictions (32).

Movement only creates potential social contacts. Whether transmission occurs or not, given a contact, is then determined by the specific transmission mechanism. For the spread of disease, transmission probabilities might depend on the route of infection, virulence and shedding levels for pathogens. For the spread of behaviours and information, transmission depends on social learning mechanisms (e.g. enhancement versus imitation) or strategies (e.g. who, what and when to learn) (33; 34; 35; 36). Transmission may be unbiased, whereby the transmission probability is proportional to the number of knowledgeable (or infected) contacts (37; 38; 39). This assumes that transmission probabilities per contact are relatively constant in time, which we refer to as “simple” transmission. Alternatively, transmission can be a more complex process. It can, for example, depend on the group composition of states amongst contacts (40; 39) or be frequency dependent. Examples of the latter include the anti-conformist transmission of song in ground finches, *Geospiza fortis*) (41), novelty biased transmission leading to revolutions in humpback whale song (*Megaptera novaeangliae*) (42; 43)), or the conformist transmission of foraging preferences in great tits (*Parus major*) (44; 45) and swamp sparrow (*Melospiza georgiana*) song (46). Complex forms of transmission are likely to alter the relationships between the patterns of contact and the dynamics of spread.

While previous studies have explored the effects of spatially explicit movement on spreading dynamics (47; 48; 49; 50; 51), these have generally not explored varying transmission mechanisms. Thus, there is an existing gap in the literature regarding the interaction between spatially explicit movement and social transmission (i.e. the social-spatial interface (52)) and its effect on the spread of behaviour. Here, we use agent-based models to test how different combinations of individual movement and transmission rules shape the emergent network structure and spreading dynamics. We borrow from the animal movement literature and use different Von Mises distributions (53) to model four spatially explicit movement rules. We also model one simple and four complex transmission rules with the widely used network-based diffusion analysis (NBDA) framework (38; 54; 55). Finally, we investigate the ability of static networks (the average of social connections over time into one network) to accurately reflect spreading dynamics of different transmission rules on dynamic networks. Together, these approaches allow us to partition out the contribution of different spatial and social behaviours to spreading dynamics, and to investigate how common methodological assumptions might impact some of the conclusions that are drawn from studies of social transmission and information spread.

## 2 Methods

We constructed a simulation model of spatially explicit populations of *N* = 196 individual agents (Figure 1A). The square number of *N* = 196 was chosen so that agents’ initial positions were evenly distributed in a lattice arrangement across the world. Agents were characterized by a position at time *t* (*x*_*it*_, *y*_*it*_), an association matrix that reflected their distance to other agents (*A*_*ijt*_), a binary knowledge state (*K*_*it*_), a movement rule (*M*), and a transmission rule (*T*). Simulations were initialized with one randomly chosen “seed” agent with knowledge of a novel behaviour. In each time-step, agents changed position according to their movement rule. They then had an opportunity to acquire the novel behaviour from their associates, defined by their association matrix and transmission rule. If they acquired a behaviour within that timestep, they updated their knowledge state. These steps were repeated until all agents were knowledgeable. From each simulation, we recorded the order of acquisition by each agent (*O*_*a*_), the time of acquisition (*T*_*a*_), and the association matrix.

**Figure 1.**
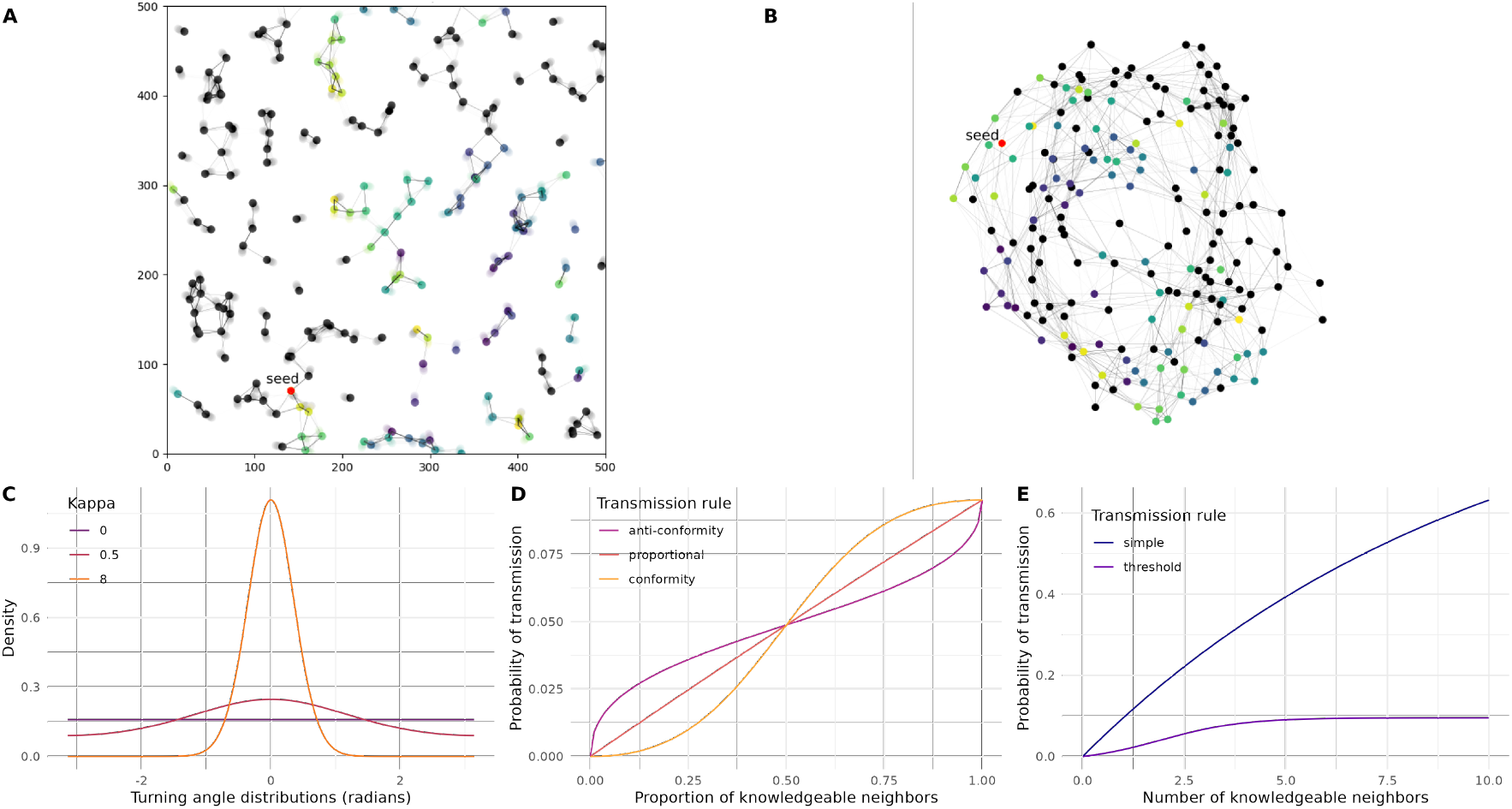
Model summary. We performed two types of simulations of spreading. **A)** In dynamic simulations, agents updated their spatial position using movement rules, and connected with other agents as they moved. At *t* = 0, one agent was randomly chosen as a seed agent (red) who knew a novel behavior, which could then spread across connections according to a transmission rule. We recorded the time agents acquired the behavior (color, black agents naive), and simulation ended when all agents acquired it. **B)** In static simulations, we created a single representation of the same network, where the edge weight represented the average connection between agents over the entire simulation. We then allowed the novel behavior to spread from the same seed agent. **C)** In dynamic simulations, we created different movement rules by varying the concentration parameter *κ* of the Von Mises distribution. For each movement rule, we tested 5 different transmission rules that considered **D)** the overall proportion of connected agents, or **E)** the number of connected agents.

Once the simulation finished, we performed a secondary simulation using the same transmission rule, except on a static representation of the network generated over the entire first simulation (Figure 1B). This allowed us to compare the performance of the transmission rule between a dynamic, “live” population, and the static network a hypothetical researcher would have measured over the course of the spread. Given the practical difficulties of recording dynamic networks in the wild, these secondary simulations provided critical insight into how spreading dynamics would play out in static representations commonly found in the literature that estimate rates of contact between individuals (see (56; 57)). In this second simulation, agents did not update their spatial position or edge weights, but rather each edge weight represented the average connection between individuals from the dynamic simulation, equivalent to a weighted simple ratio index (SRI; equation 5). The seed agent was also identical to the first simulation to keep initial conditions of the spread the same.

### 2.1 Movement rules

We tested four movement rules that resembled various strategies ranging from completely random, localized movement to purpose-driven movement. Agents were each initialized with a random starting direction and moved on a 2d toroidal surface with an area of 500 units squared. Using toroidal geometry ensured that there were no boundaries to the environment, as x and y coordinates wrapped around each other. Agents moved at a fixed velocity of 1 unit per timestep while choosing turning angles drawn from a circular-normal Von Mises distribution, similar to widely used step-selection models (53). The Von Mises distribution is characterized by two parameters, mean direction *µ* and concentration parameter *κ*. We created various movement rules by changing *κ*, which affected the spread of turning angles.

1. **Localized movement** – Agents moved in a random direction during each timestep. For each timestep, a direction was chosen from a von Mises distribution where concentration parameter *κ* = 0, equivalent to a uniform distribution of angles between 0 and 2*π*.
2. **Semi-localized movement** – Agents largely moved in the same area, but occasionally would deviate away from that area. For each timestep, the direction was chosen from a von Mises distribution where *κ* = 0.5.
3. **Nomadic movement** – Agents largely moved straight paths with very occasional deviations. Each timestep, the direction was chosen from a von Mises distribution where *κ* = 8.
4. **Resource-driven movement** – Agents navigated toward or away from 9 resources arranged in a grid based on their satiety state, which depended on their proximity to resources. If the agent was “hungry”, it moved toward the closest resource. Agents updated their satiety state to “full” when they reached a resource and then moved away from the resource for 50 timesteps. Turning angles were chosen based on a Von Mises distribution where *µ* = *±*resource direction and *κ* = 8, resulting in a relatively straight path similar to the nomadic rule.

For rules 1 through 3, *µ* could change over time, as the angle chosen at time *t* became *µ* at time *t* + 1. For rule 4, *µ* was not recursive and instead depended on the resource location and state. We have provided visualizations of exemplar simulations in supplementary videos S1-S4 to provide the reader with an intuition of each movement rule.

### 2.2 Edge weights

These movement rules led to the emergence of a network structure that changed with each timestep. Edge weight *A*_*ijt*_ between agents *i* and *j* at time *t* were based on the proximity of agents. We calculated edge weights by first computing the toroidal distance *d* at time *t* between agents *i* and *j* with positions *x*_*it*_, *y*_*it*_ and *x*_*jt*_, *y*_*jt*_ on the 500 units squared toroidal surface:

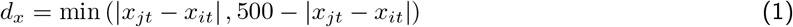

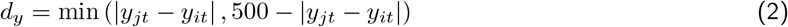

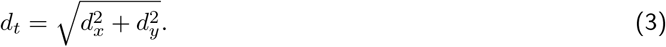

Once *d*_*t*_ was calculated, we used a threshold to only consider agents as neighbours within 34 units of the agent using

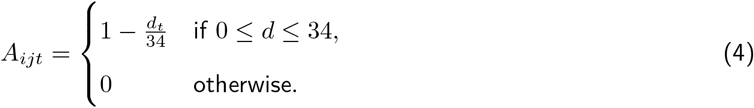

The weight decreased linearly with distance and became zero if the distance was greater than the threshold. To reduce simulation variance in order to isolate the effect of movement rules, agents were initialized as equally spaced out on the toroidal grid. The threshold of 34 units was calculated such that no agents were connected upon initialization, as the initial distance between agents was *≈* 36 units. Thus, all subsequent associations were formed solely on the movement rules, rather than chance initial conditions.

Once the simulation finished running, we created a static representation of the dynamic network by summing edge weights and then dividing by the timesteps the simulation took to finish running. Thus, the static association matrix *A*_*ij*_ had edge weights defined as:

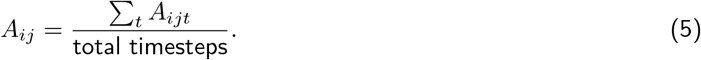

This equation is equivalent to the simple ratio index, a commonly used measure of association in animal social networks (57). Given that we have complete information about the population, SRI terms that consider periods when only *i* or *j* are observed are 0, leaving the denominator as only the total sampling periods. The only difference is that we have summed the weighted measure of association for each sampling period in the numerator, rather than a binary measure.

### 2.3 Transmission rules

At each timestep, naive agents had the opportunity to learn a novel behaviour from knowledgeable associates. The transmission rules *T* were a function of these edge weights encoded in an association matrix *A*, and a knowledge state vector *K*. Transmission rules directly influenced the probability of acquiring the novel behaviour *Pr*(acquire) by naive agent *i* at time *t*. To calculate these probabilities, we use a simplified dynamic described by the general transmission model of network-based diffusion analysis (55; 54), and first calculated a rate of transmission *λ*_*i*_(*t*) as

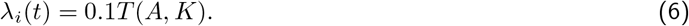

In this equation, 0.1 is a scaling factor that can be interpreted as how easy a behaviour is to be socially transmitted, or an individual’s propensity for social learning. This could take any positive value. We chose 0.1, as larger values raised the probability of transmission to unrealistically high levels and erased differences between transmission rules. After the transmission rate was calculated, this was converted into a probability *Pr*(acquire) by

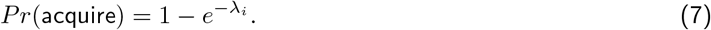

For each agent, we first calculated two sums, *n* =∑_*j*_ *A*_*ijt*_*K*_*jt*_ and *m* = ∑_*j*_ *A*_*ijt*_(1 *− K*_*jt*_), where *n* was the sum of weights of knowledgeable neighbours and *m* was the sum of weights of naive neighbours at timestep *t*. We tested one simple transmission rule and four complex transmission rules adapted from (58):

1. **Simple transmission rule**: The acquisition rate was proportional to the sum of the weights of knowledgeable neighbours:

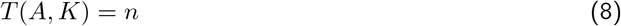
2. **Anti-conformity rule**: The probability of transmission changed non-linearly with the proportion of connected knowledgeable and naive states. Here agents were more likely to acquire a behaviour if a minority of their associates were knowledgeable.

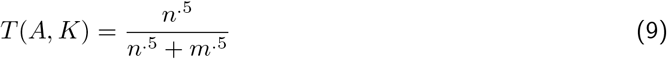
3. **Proportional rule**: The acquisition rate was proportional to the sum of the weights of knowledgeable neighbours divided by the sum of weights for all neighbours:

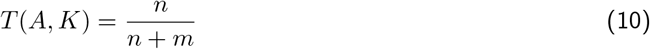
4. **Conformity rule**: Agents were more likely to acquire a behaviour if a majority of their associates were knowledgeable.

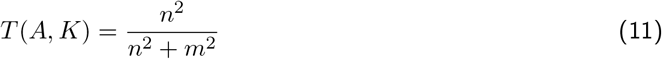
5. **Threshold rule**: A logistic function was used to calculate the probability of transmission, where agents had a threshold level *α* of knowledgeable neighbours they needed to acquire the behaviour. A sharpness parameter *s* controlled the steepness of the logistic curve. We set *α* = 2 and *s* = 1.

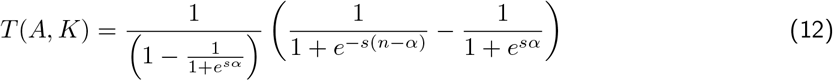

After calculating the probability of acquisition based on the selected rule, a random number was generated. If this number was less than the calculated acquisition rate, the agent acquired the behaviour, turning from naive *K*_*i*_ = 0 to knowledgeable *K*_*i*_ = 1.

### 2.4 Measurements and conditions

For each simulation, we recorded the global time-to-spread (TTS), defined as the time it took for all agents to become knowledgeable measured in timesteps. We also recorded the order of acquisition and timestep of acquisition for each agent. From this data, we could also calculate the rate of acquisitions. Once the dynamic simulation finished, we took its static representation of the network and re-ran the simulation with the same seed agent and transmission rule, but without the sequence of movement that generated the network. We ran N=100 dynamic simulations and N=100 static simulations for each combination of movement rules and transmission rules.

In order to assess how network structure might affect transmission rates, we calculated key metrics for each network: average degree, average weighted degree, average weighted clustering coefficient, and average effective distance. We calculated both average degree and average weighted degree since average weighted degree alone could not disambiguate cases where agents were weakly connected to many agents, or strongly connected to few agents. Average degree (*D*) was defined as:

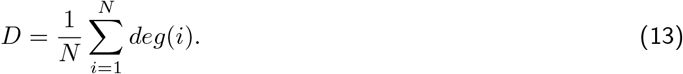

Average weighted degree (*D*_*𝓌*_) was defined as

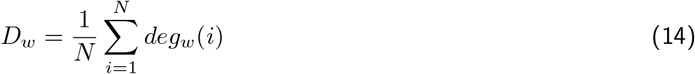

where the degree of each node *deg*_*𝓌*_ (*i*) was the sum of its weighted edges.

The average weighted clustering coefficient (*C*_*𝓌*_) measured the degree to which agents tended to form connected triangles, and was defined as

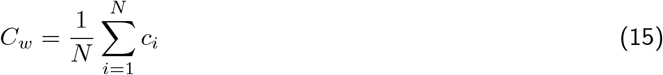

where *c*_*i*_ was the weighted clustering coefficient of agent *i* was calculated as in Barrat et al. (59):

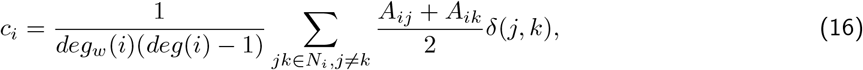

where *N*_*i*_ was the set of all neighbours connected to focal agent *i*, and δ (*j, k*) = 1 if there was an edge between neighbours *j* and *k*, else 0. If *N*_*i*_ *<* 2, then *c*_*i*_ = 0.

We used average effective distance (*E*) to account for the topological structure of the network and weights between edges. This measure is more appropriate than path length when edge weights represent costs of information flow, such as in our case where distance negatively affected transmission. We first took the negative log of edge weights by

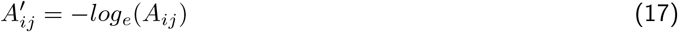

assuming 0 *≤ A*_*ij*_ *≤* 1. We then calculated the effective distance *e* as the shortest path length between *i* and *j* accounting for *A*′:

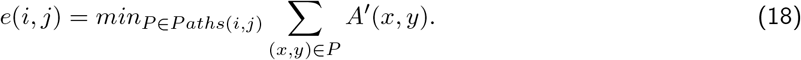

Finally, we then took the average of these effective distances *E*:

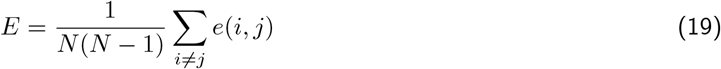

Finally, we quantified how fragmented networks were by counting the number of isolated components (including single agents), as well as the median size of all components.

In order to understand the networks generated by each movement rule, we conducted 10 preliminary simulations of 1000 timesteps for each movement rule. These were performed separately from the main transmission simulations described above. We recorded these key network metrics both cumulatively over the course of the simulation and instantaneously at each timestep.

We report variation for our recorded metrics using both the coefficient of variation (CV), and the percentile interval (PI), which describes the interval of a distribution that contains 95% of variation, with equal weight assigned to each tail.

We note that we have not performed an exhaustive sensitivity analysis of all possible parameters, and instead have created reasonable characterizations of movement and transmission rules that distinct, but not extreme. With this study, we aimed to ask 1) whether spreading dynamics of transmission mechanisms differ depending on movement rules, and 2) whether static representations of dynamic networks predict the same spreading dynamics. The precise effects of intermediate or edge-case parameterizations are not critical to this demonstration.

## Results

### 3.1 Network structure depended on movement rules

Our four movement rules produced different global structures (visualized in Figure 2A), with divergent patterns found between cumulative and instantaneous metrics (Figure 2B-G). Overall, we found very little simulation variance in these network metrics, as evidenced by the tight percentile intervals in Figure 2. We first explored measures of degree, average degree *D* (Figure 2B,C) and average weighted degree *D*_*𝓌*_ (Figure 2D,E), as we expected more connected networks to increase transmission rates throughout the simulation, especially for simple transmission. There were significant differences in the cumulative metric (Figure 2B,D)—the aggregation of all observed data up to that time point into a static network—of network connectivity as the simulation progressed. Localized movement obtained the lowest *D* and *D*_*𝓌*_, indicating that agents were weakly connected to few agents. Nomadic movement resulted in the highest *D*, yet obtained a similar *D*_*𝓌*_ to semi-localized movement. Thus, nomadic individuals were weakly connected to many agents, while semi-localized agents were more strongly connected to fewer agents. Resource-based movement obtained a lower *D* than semi-localized movement, yet obtained the highest *D*_*𝓌*_, indicating that agents were even more strongly connected to fewer agents.

**Figure 2.**
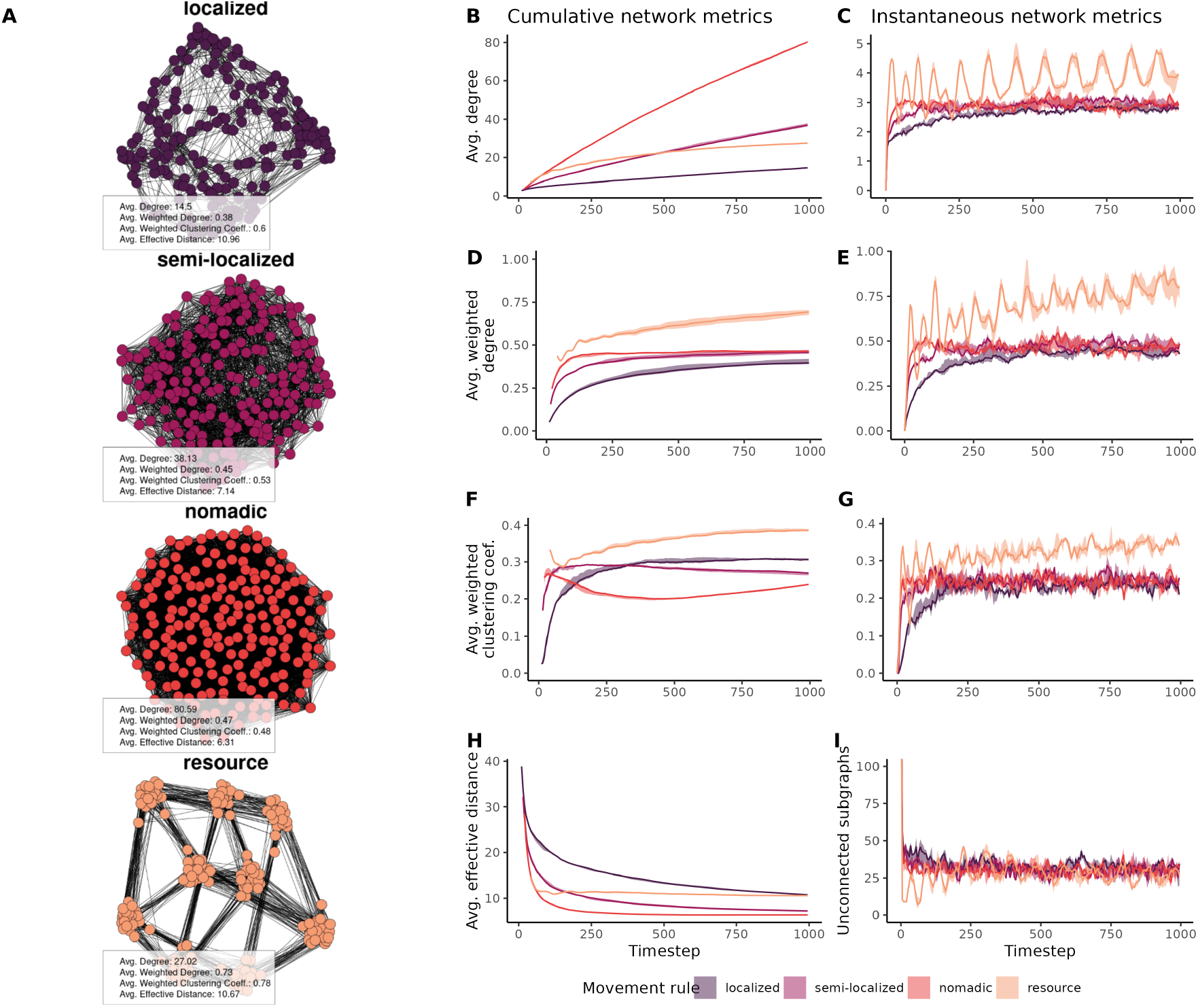
Movement rules generate different network structures. **A)** Visualization of exemplar static networks generated by four different movement rules, with edge data recorded for 1000 timesteps (N=10 simulations per rule). **B-D)** Mean (center line) and 95% PI (shade) of cumulative network metrics under different movement rules. **E-G)** Mean and 95%PI of instantaneous network metrics generated by the same simulations.

Interestingly, this pattern was quite different when looking at network structure for each timestep independently (instantaneous metrics, Figure 2C,E). Localized, semi-localized, and nomadic movement all resulted in similar instantaneous *D* and *D*_*𝓌*_, although localized movement was still lower than the others. Resource-based movement resulted in significantly higher *D* and *D*_*𝓌*_ that oscillated as agents moved towards and away from resources.

We next explored the average weighted clustering coefficient *C*_*𝓌*_, which indicated the degree to which individuals tended to form connected triangles. When looking at the cumulative metrics, we found that resource-based movement resulted in the highest *C*_*𝓌*_ by the end of the simulation, followed by localized, semi-localized and finally nomadic movement (Figure 2F). Instantaneous metrics followed a similar pattern to degree metrics, where resource movement resulted in a slight oscillation around 0.3, and all other rules oscillated around a slightly lower value (Figure 2G).

Finally, we explored average effective distance *E* on cumulative networks (Figure 2H), which represented the ease with which information could travel from one individual to any other individual in the network via the shortest possible weighted path. We found that effective distance all decreased over time, with localized and resource movement rules obtaining similar longest effective distances at *E* = 10.96 and *E* = 10.67. However, effective distance decreased at a much slower rate with localized movement. Semi-localized and nomadic movement obtained shorter distances, at *E* = 7.14 and *E* = 6.31. As effective distance could not be computed when the network was fragmented, we could not explore *E* for instantaneous networks. Instead, we explored the number of isolated components (Figure 2I). Interestingly, we found that all four rules resulted in a similar number of isolated components across the entire simulation.

We expected that these differences between instantaneous and cumulative metrics might contribute to variable performance of our transmission rules when tested *in situ* during dynamic network simulations, or on the static, cumulative representation of that same network, which we go on to test in the next two sections.

### 3.2 Transmission rule performance depended on movement rules

We next compared the performance, in terms of time-to-spread (TTS), of our transmission rules. When comparing the relative ranks of transmission rules within each movement rule, we found a consistent pattern in dynamic networks: under all movement rules except resource, anti-conformity always spread the fastest, followed by proportional, conformity, simple transmission and finally threshold rules (Figure 3, dynamic networks panel; for specific values and coefficients of variation see Table S1). Interestingly, under resource-based movement, simple transmission outperformed conformity. This pattern was likely due to the high clustering and average weighted degree of resource-based movement created larger probabilities for transmission under the non-normalized term *T* (*A, Z*) under simple transmission, compared to the normalized *T* (*A, Z*) under conformity. Furthermore, the conformity rule would result in slower initial spread of information within a given cluster of individuals.

**Figure 3.**
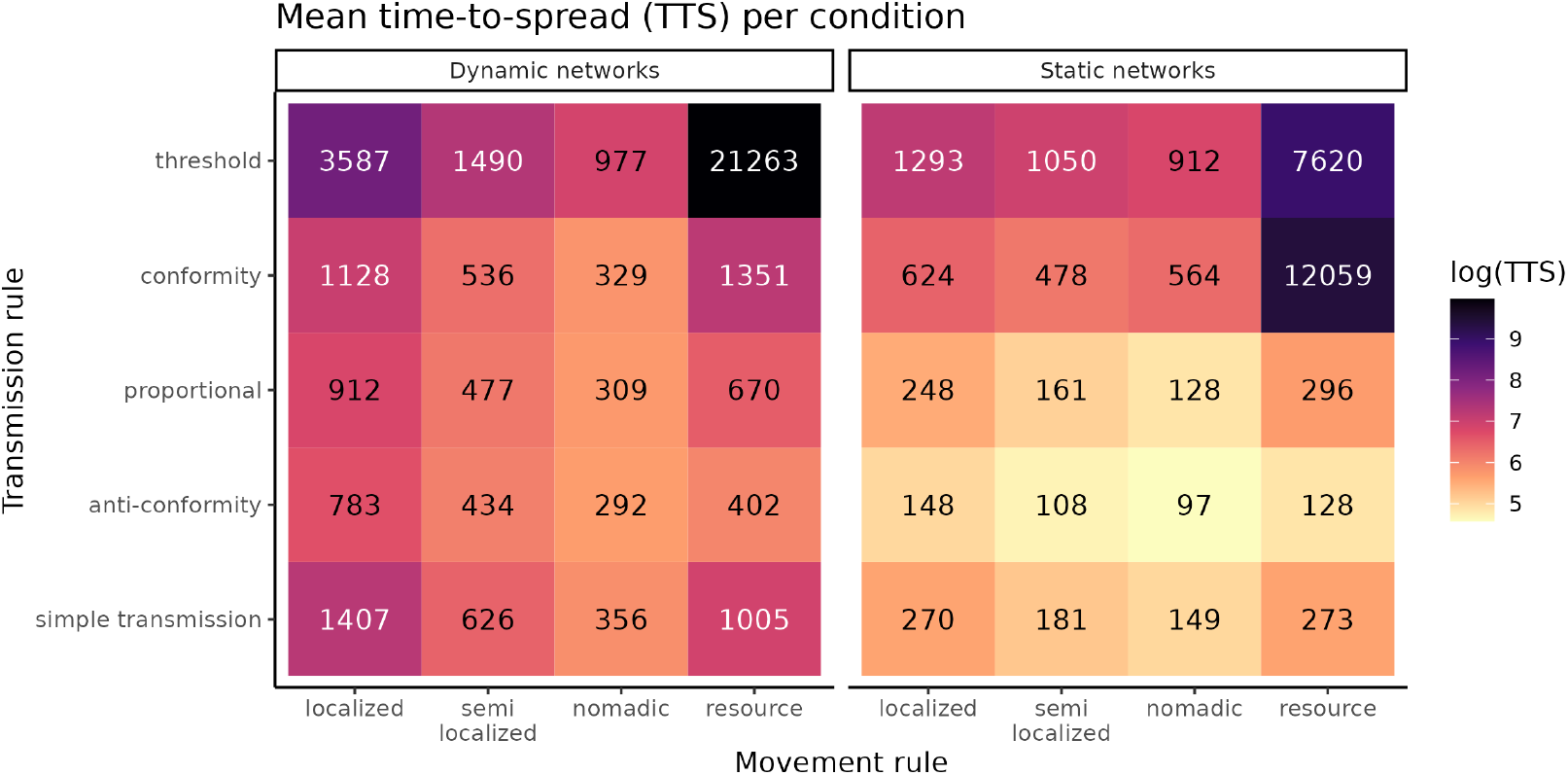
Transmission rules vary in performance depending on movement rules. Mean time-to-spread for each combination of movement rule (x-axis) and transmission rule (y-axis).

When comparing movement rules within each transmission rule, nomadic movement rules always resulted in the quickest spread Figure 4. However, the relative ranks of the remaining movement rules depended on the transmission rule. Under simple transmission and proportional rules, nomadic was followed by semi-localized, resource, and finally localized obtaining the slowest spreads. Anti-conformity obtained slightly slower spread under resource-based movement, then semi-localized and finally localized. Conformity obtained a ranking of nomadic, semi-localized, localized and resource. The threshold rule showed the most variation, with resource-based movement spreading two orders of magnitude slower than the best-performing combination of anti-conformity and nomadic movement, and an order of magnitude slower than localized movement. When examining the spreading rate over time, this appeared to be caused by 1) a slow start at the beginning of simulations, and 2) a strong deceleration in transmission rates at the end of the simulation. This deceleration was likely caused by the routine movement of agents motivated by resource locations. The probability of reaching the threshold value for transmission into clusters of uninformed individuals was reduced to the stochastic immigration of enough agents from other resource patches, which could take thousands of timesteps.

**Figure 4.**
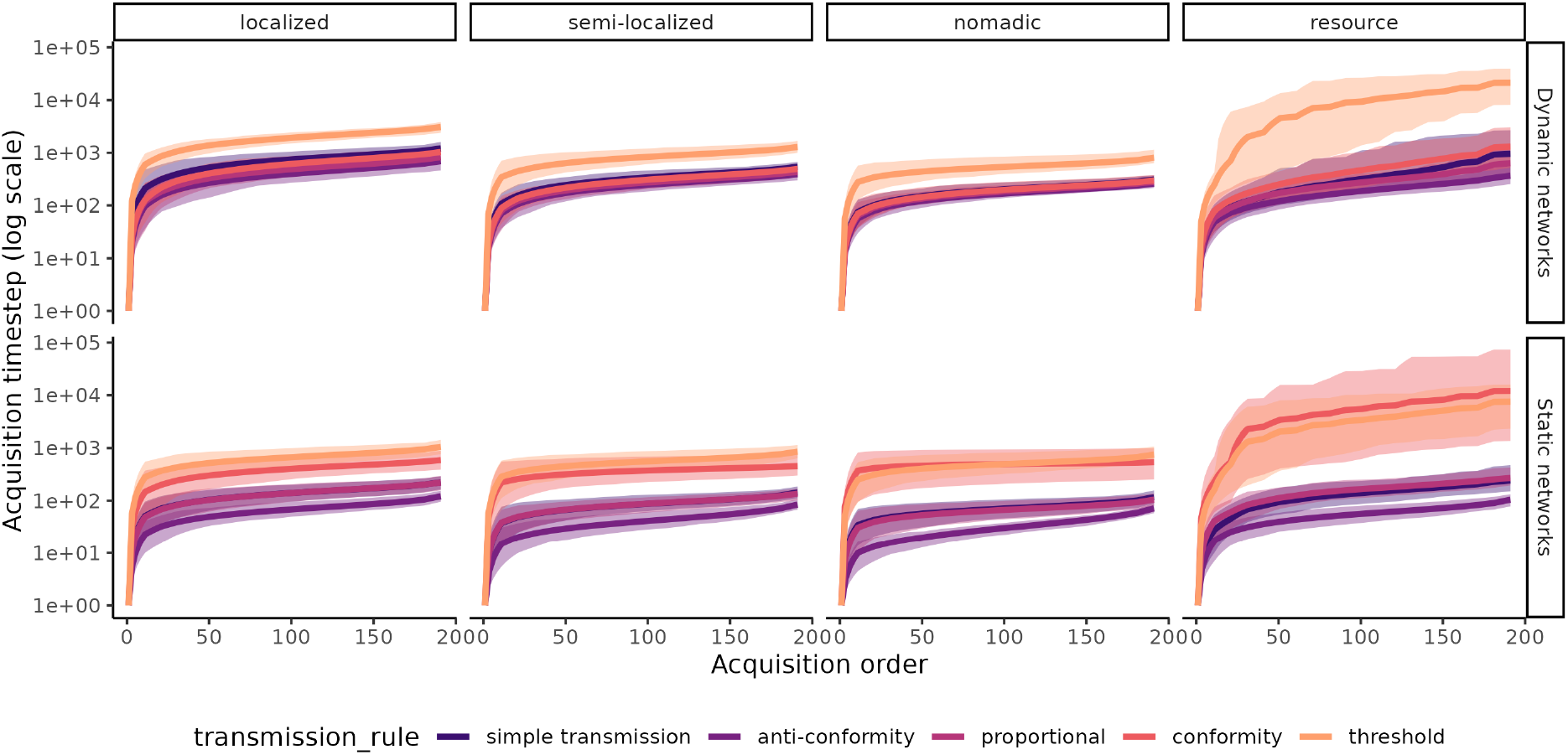
Spreading dynamics between dynamic and static network representations. Order of acquisition (x-axis) and timestep of acquisition (y-axis, log for readability). Mean (center line) with 95% PI (shade) shown for N=100 simulations for each combination of conditions.

### 3.3 Transmission rules performed differently when tested on static representations of dynamic networks

We examined spreading dynamics when re-running simulations with the same seed agent on a static representation of the network taken cumulatively throughout the simulation. We largely found that *TTS* was lower than in the dynamic network simulations, with several notable changes in the relative rankings (Figure 3, static networks panel; Table S1 for specific values). Firstly, when comparing different movement rules under simple transmission, both localized and resource-based movement obtained nearly identical *TTS*, whereas under dynamic networks, behaviours spread significantly slower under localized movement, compared to resource-based movement. Next, when comparing transmission rules under resource-based movement, conformity was slower than threshold by an order of magnitude, becoming the slowest combination overall. Finally, the extremely slow spread of threshold transmission under resource movement was reduced by an order of magnitude, although the relative ranking of movement rules within the threshold rule remained the same.

There were also notable differences between static and dynamic networks regarding the magnitude and timing of changes to the rate of spread over time, visualized in Figure 5. Static representations of movement resulted in exaggerated differences in spreading rates between all rules compared to the dynamic networks. Simple transmission and proportional rules resulted in similar acceleration patterns, although proportional tended to reach its peak spreading rate slightly after simple transmission. Spreading rates for anti-conformity rules reached a dramatic maximum much earlier in simulations, although markedly decelerated into the final acquisition events. However, this quick start still enabled anti-conformity to outperform other rules. Interestingly, the conformity rule resulted in an acceleration towards the end of simulations in static network simulations, but not dynamic network simulations. This acceleration was most pronounced under nomadic movement. Both threshold and conformity were generally much slower in the initial stages of spreading compared to other rules. Under resource movement, the conformity rule evidenced a much slower spread early on, which was not found in the dynamic networks. We believe that this resulted from the lack of spatially explicit positions of agents in the static representation of the network. The behaviour had difficulty spreading within the cluster of agents which held the seed agent (e.g. one of the clusters shown in Figure 2A, resource), and had further difficulty in spreading to any other cluster. When agents were spatially explicit, they might frequently be in smaller sub-graphs, which benefited the normalized *T* (*A, K*) of conformity, allowing for the behaviour to spread more easily.

**Figure 5.**
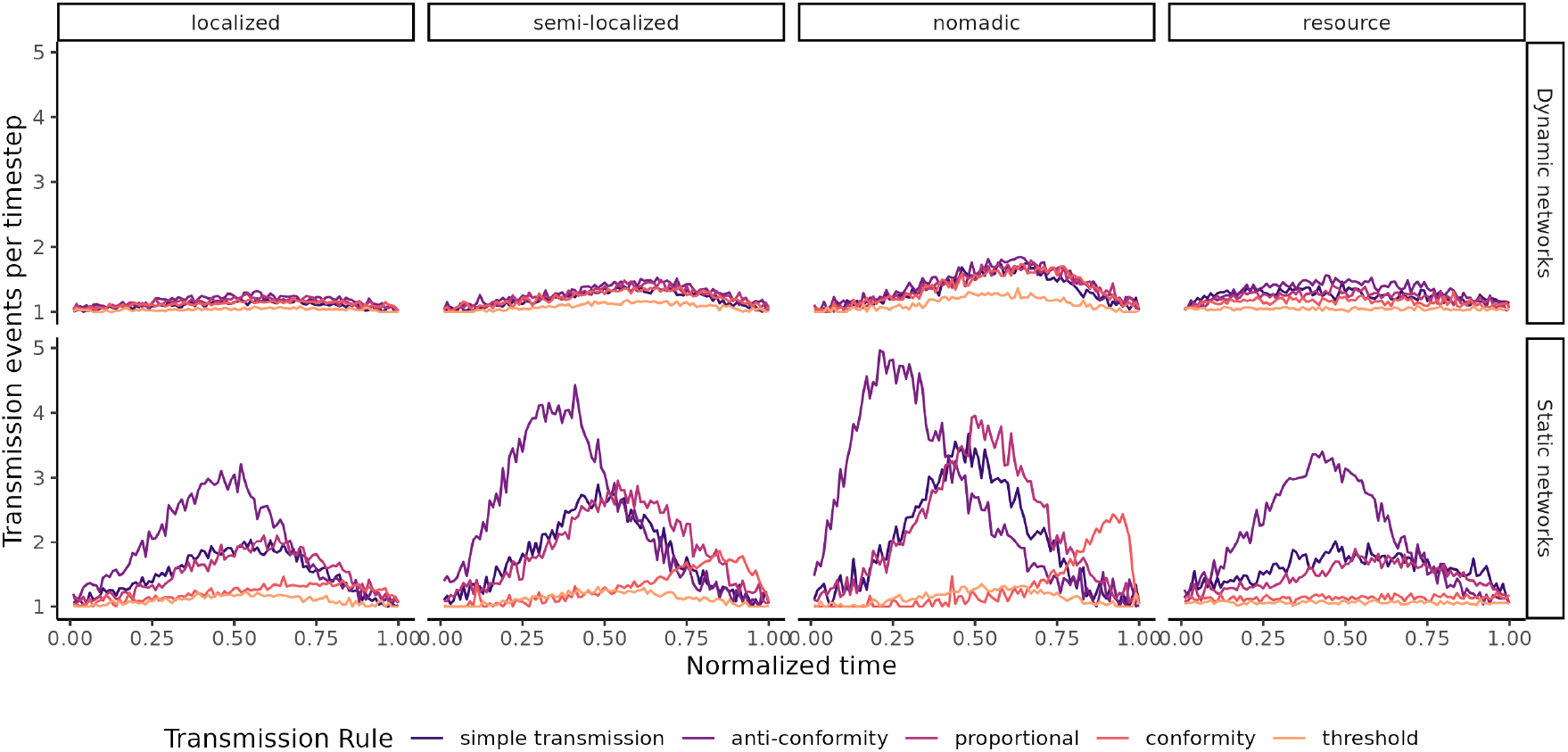
Spreading rates over normalized timesteps. Mean rate of new acquisitions over time under different movement and transmission rules. Variation measurement not included for readability. Time was normalized for each simulation by dividing by the timestep of the final acquisition event. The rate of spread over time differed in magnitude and pattern over time between dynamic and static networks.

## 4 Discussion

We have shown that different movement rules can generate substantial differences in the properties of the resulting social networks. These differences then have consequences for spreading dynamics. However, the translation of movement rules to spreading dynamics—such as which movement rules produce the fastest spread to all individuals—depended on the transmission rules that individuals use. Furthermore, we have demonstrated that predictions about spreading dynamics can differ in rate and acceleration between dynamic networks and static representations of those networks. These findings, therefore, highlight how spatial and social processes can interact to shape ecological (what traits individuals express) and evolutionary (how behavioural traits affect fitness) feedbacks.

Importantly, we found that the spreading dynamics of both simple and complex transmission rules changed between dynamic networks, and static representations of those networks. For example, the transmission rate was higher for simple transmission in static networks, resulting in much faster spreads, and conformity evidenced a strong acceleration in static networks towards the end of the spread, which was not found in dynamic networks. Such discrepancies, as well as differences in cumulative and instantaneous network metrics, are relevant for empirical researchers, who usually construct cumulative networks over several weeks, months, or years of data collection (60; 11; 61; 62; 10; 63). The implication of our findings is that our ability to faithfully capture the relative contribution of movement and transmission rules to spreading dynamics is likely to be affected by methodological decisions and limitations in the study design. Specifically, studying spread on aggregated networks–those that average the rates of contact among individuals over time–can produce different outcomes than studying the same behaviours in the underlying dynamic network (even when the static network faithfully captures the complete patterns of contact). These results extend previous work finding that static networks only provide meaningful representations of social connections when the relationship between the connection strength and the probability of transmission taking place is linear (32). We suggest instead that studies should aim to faithfully capture the temporal patterning of the study population when studying spreading dynamics on networks. A further finding of our study is that accounting for the contribution of movement to network dynamics is also important. Prior models of spreading on dynamic social networks abstracted the contribution of movement rules to changes in networks (29; 31; 27; 6; 64), whereas here we have explicitly modelled these decisions using a 2D coordinate system. Spatially explicit movement explains some interesting differences between our findings and previous work. For example, Springer et al. (31) found that dynamic networks generally resulted in larger outbreaks, which suggested a higher overall transmission rate (although they did not include a comparable measure to our time-to-spread). In contrast, we found that spreading happened significantly faster on static representations of dynamic networks. This indicates that implicitly accounting for movement in dynamic networks, without modelling the time it takes agents to actually reposition themselves and thus rewire their connections, does not allow for accurate estimates of the rate of spreading. One reason for this is that there may be some temporal correlations in contact rates among individuals that cannot be easily replicated in space-free models. Specifically, if individuals A and B are in contact, and then C comes into contact with A, then C will be more likely to also come into contact with B than it will be most other individuals in the population—simply because A, B, and C are all sharing the same local spatial area. This finding highlights the importance of the spatial-social interface in shaping spreading dynamics.

While our study touched on the effect of state-based movement patterns with our resource rule, we did not explore movement that depended on social cues or knowledge state (e.g. move towards knowledgeable agents). Preferential avoidance and attraction towards conspecifics are important behaviours in real-world social systems, evolving as a means to maximize fitness. For example, human and non-human animals will preferentially avoid infected conspecifics (7; 5). In the context of information transmission, a recent study on the co-evolution between social networks and cultural transmission demonstrated how a trade-off between skill generalization and specialization could result in sparser or denser networks (65). While spreading rate was not the focus of that study, it would be an interesting to explore both 1) how transmission could lead to changes in social networks, and 2) how spreading dynamics might change based on social state-based movement rules. For example, if individuals preferentially follow knowledgeable agents, networks would likely organize into a scale-free topology (66). Under such conditions, we would also expect faster spreading, however this may be tempered by factors such as competition and finite resources (67). For example, increasing aggregation sizes following information transmission might drive behaviours that prevent information spread, such as monopolization of resources by dominant individuals or strategic use of information to prevent others from observing the information. Furthermore, if a resource is finite, once the resource is depleted then opportunities for the transmission of the knowledge required to exploit that resource would also vanish (68).

Our results may also be compared and contrasted with models of complex transmission in the context of consensus. Consensus models are subtly different in that agents can flexibly switch opinion states many times depending on neighbours’ states, rather than the one-way change in our model from naive to knowledgeable. A prior study found that any network structure reached consensus slower than well-mixed (complete) networks (69), which matched our result that nomadic movement obtained the fastest spread, independent of transmission rule. The study further revealed that modular networks could greatly increase the time to consensus. Our most modular network was generated by resource-based movement, and this resulted in a very long time-to-spread under the threshold rule. However, other rules resulted in a slower spread with localized movement. The spreading dynamics of conformity, in particular, depended on whether networks were dynamic or static representations. Therefore, it may be fruitful to revisit the context of consensus with a comparison between dynamic and static representations of networks.

In summary, our study has highlighted interactions at the interface between spatial movement rules and social transmission rules that resulted in variable performance in the spread of information. Our results also emphasize that predictions about transmission might change depending on how networks are represented, especially when transmission is not linearly related to connectivity (i.e. complex transmission). Future studies may build on these findings to further elucidate the complicated feedback loop between culture, movement and social network structure.

## 5 Data Availability

Code and data for statistical analyses and main text figures are publically available on Edmond at https://doi.org/10.17617/3.U3NSZT (70).

## 6 Acknowledgements

This work was supported by the Max Planck Society and the Centre for the Advanced Study of Collective Behaviour, funded by the Deutsche Forschungsgemeinschaft (DFG) under Germany’s Excellence Strategy (EXC 2117-422037984). DRF was funded by the European Research Council (ERC) under the European Union’s Horizon 2020 research and innovation programme (No. 850859) and an Eccellenza Professorship Grant of the Swiss National Science Foundation (PCEFP3 187058).

## 7 Author Contributions

Conceptualisation, M.C., D.R.F.; Methodology, M.C., D.R.F.; Software, M.C.; Investigation, M.C.; Visualization, M.C.; Writing - Original Draft, M.C., D.R.F.; Writing - Review & Editing, M.C., D.R.F.

## 8 Competing Interests

The authors declare no competing interests, financial or otherwise.

## Notes

https://doi.org/10.17617/3.U3NSZT

### Competing Interest Statement

The authors have declared no competing interest.

### Summary of Updates

Added summary figure for model. Light edits for clarity and concision.

