## Supplementary material for "The contribution of movement to social network structure and spreading dynamics under simple and complex transmission": table S1

### Tables

- **Table S1**: Summary statistics for time-to-spread.

### Videos

- **Video S1**: Exemplar simulation with localized movement rule and proportional transmission rule.
- **Video S2**: Exemplar simulation with semi-localized movement rule and proportional transmission rule.
- **Video S3**: Exemplar simulation with nomadic movement rule and proportional transmission rule.
- **Video S4**: Exemplar simulation with resource based movement rule and proportional transmission rule.

Table S1: **Summary statistics for spreading dynamics.** Mean time-to-spread (TTD), coefficient of variation (CV), min and max 95% PI for both *in situ* simulations on dynamic networks, and simulations using static representations of those networks.

| | Dynamic networks | | | | Static networks | | | |
|---|---|---|---|---|---|---|---|---|
| movement rule | mean TTD | CV | PI min | PI max | mean TTD | CV | PI min | PI max |
| **simple transmission** | | | | | | | | |
| localized | 1406.51 | 14.95 | 1139.57 | 1752.11 | 270.22 | 15.50 | 204.45 | 332.11 |
| semi-localized | 626.36 | 13.45 | 503.34 | 773.00 | 181.24 | 17.31 | 142.89 | 240.56 |
| nomadic | 355.83 | 10.22 | 304.34 | 417.11 | 149.30 | 17.79 | 109.44 | 196.11 |
| resource | 1004.89 | 76.67 | 438.89 | 2247.58 | 273.01 | 27.94 | 189.12 | 407.66 |
| **anticonformity** | | | | | | | | |
| localized | 783.02 | 16.65 | 569.36 | 996.11 | 148.49 | 13.06 | 120.34 | 178.56 |
| semi-localized | 433.94 | 14.66 | 353.00 | 548.55 | 108.26 | 13.22 | 88.34 | 128.00 |
| nomadic | 292.46 | 9.76 | 251.00 | 333.00 | 96.88 | 16.73 | 77.44 | 127.11 |
| resource | 401.53 | 22.07 | 287.89 | 548.33 | 127.74 | 15.84 | 95.44 | 158.67 |
| **proportional** | | | | | | | | |
| localized | 911.56 | 16.30 | 709.12 | 1148.77 | 248.22 | 15.22 | 184.89 | 313.11 |
| semi-localized | 477.25 | 10.19 | 413.00 | 563.88 | 160.82 | 16.02 | 129.44 | 186.67 |
| nomadic | 308.57 | 10.05 | 272.44 | 376.22 | 128.10 | 15.16 | 101.00 | 159.56 |
| resource | 669.84 | 52.53 | 384.00 | 1183.51 | 296.14 | 22.29 | 221.44 | 410.55 |
| **conformity** | | | | | | | | |
| localized | 1128.08 | 15.79 | 848.68 | 1391.88 | 623.85 | 22.72 | 432.78 | 856.11 |
| semi-localized | 536.47 | 10.87 | 453.34 | 626.78 | 478.40 | 30.84 | 330.35 | 713.66 |
| nomadic | 328.66 | 10.80 | 282.34 | 398.78 | 563.74 | 35.19 | 308.46 | 967.11 |
| resource | 1350.67 | 51.83 | 585.89 | 2729.30 | 12059.33 | 169.98 | 1480.57 | 39672.25 |
| **threshold** | | | | | | | | |
| localized | 3586.94 | 12.82 | 2911.93 | 4294.71 | 1293.04 | 17.23 | 942.67 | 1734.75 |
| semi-localized | 1490.38 | 14.54 | 1206.80 | 1869.22 | 1050.31 | 13.95 | 860.22 | 1301.54 |
| nomadic | 976.55 | 15.81 | 774.56 | 1259.53 | 912.34 | 15.70 | 703.90 | 1152.54 |
| resource | 21263.36 | 39.09 | 8885.35 | 36108.72 | 7620.02 | 51.92 | 2760.82 | 14291.54 |
